# Using marine cargo traffic to identify countries in Africa with greatest risk of invasion by *Anopheles stephensi*

**DOI:** 10.1101/2021.12.07.471444

**Authors:** Jordan Ahn, Marianne Sinka, Seth Irish, Sarah Zohdy

## Abstract

*Anopheles stephensi* is an efficient malaria vector commonly found in South Asia and the Arabian Peninsula, but in recent years it has established as an invasive species in the Horn of Africa (HoA). In this region, *An. stephensi* was first detected in a livestock quarantine station near a major seaport in Djibouti in 2012, in Ethiopia in 2016, in Sudan in 2018 and Somalia in 2019. *Anopheles stephensi* often uses artificial containers as larval habitats, which may facilitate introduction through maritime trade as has been seen with other invasive container breeding mosquitoes. If *An. stephensi* is being introduced through maritime traffic, prioritization exercises are needed to identify locations at greatest risk of *An. stephensi* introduction for early detection and rapid response, limiting further invasion opportunities. Here, we use UNCTAD maritime trade data to 1) identify coastal African countries which were most highly connected to select *An. stephensi* endemic countries in 2011, prior to initial detection in Africa, 2) develop a ranked prioritization list of countries based on likelihood of *An. stephensi* introduction for 2016 and 2020 based on maritime trade alone and maritime trade and habitat suitability, and 3) use network analysis to describe intracontinental maritime trade and eigenvector centrality to determine likely paths of further introduction on the continent if *An. stephensi* is detected in a new location. Our results show that in 2011, Sudan and Djibouti were ranked as the top two countries with likelihood of *An. stephensi* introduction based on maritime trade alone, and these were indeed the first two coastal countries in the HoA where *An. stephensi* was detected. Trade data from 2020 with Djibouti and Sudan included as source populations identify Egypt, Kenya, Mauritius, Tanzania, and Morocco as the top five countries with likelihood of *An. stephensi* introduction. When factoring in habitat suitability, Egypt, Kenya, Tanzania, Morocco, and Libya are ranked highest. Network analysis revealed that the countries with the highest eigenvector centrality scores, and therefore highest degrees of connectivity with other coastal African nations were South Africa (0.175), Mauritius (0.159), Ghana (0.159), Togo (0.157), and Morocco (0.044) and therefore detection of *An. stephensi* in any one of these locations has a higher potential to cascade further across the continent via maritime trade than those with lower eigenvector centrality scores. Taken together, these data could serve as tools to prioritize efforts for *An. stephensi* surveillance and control in Africa. Surveillance in seaports of countries at greatest risk of introduction may serve as an early warning system for the detection of *An. stephensi*, providing opportunities to limit further introduction and expansion of this invasive malaria vector in Africa.

## Introduction

Globalization and the movement of humans and goods has facilitated the introduction of organisms to new locations, and the list of invasive species has grown substantially since the 1980s (Seebens et al., 2020). From 2006 to 2014, the movement of maritime shipping between socio-economic regions, defined as maritime countries grouped by similar socio-economic factors, increased by 258% with projected growth of maritime movement of 240% to 1,209% from 2014 to 2050 (Sardain et al., 2019). However, invasive species are not limited to organisms like zebra mussels (Haag, K. H., 1994), pine and eucalyptus trees (Global Invasive Species Database, 2021; Ritter & Yost, 2009), and feral hogs (*USDA APHIS* | *History of Feral Swine in the Americas*., n.d.). Invasive species can also include arthropod vectors of disease and microbial agents, posing significant public health threats. A prime example is the introduction of *Aedes aegypti*, the yellow fever mosquito, through the movement of ships in the 19^th^ century to the Americas (Powell et al., 2018). In the 20^th^ century, further movement of cargo ships, in particular those carrying used tires, facilitated the spread of *Aedes* spp., including *Aedes albopictus*, a successful invasive species, which is now established on six continents. The proposed mechanisms facilitating the success of *Aedes* spp. invasion via ships include a few characteristics common to *Aedes aegypti* mosquitoes: the use of artificial containers as larval habitats, the preference for human blood meals, and the ability for eggs to resist desiccation in the absence of water. This drought tolerance has been proposed as a key explanation for why *Aedes* species spread efficiently by sea, as less drought-tolerant species may require more rapid transportation, such as air travel, for invasive populations to survive long enough to establish in a new location. Other mosquitoes, such as *Anopheles* mosquitoes, have also created great public health challenges when accidentally introduced to non-native countries such as Egypt and Brazil. *Anopheles arabiensis* was the cause of malaria outbreaks in Brazil, but was eventually eradicated after challenging and well-coordinated control measures were put in place (Killeen et al., 2002).

Unlike *An. arabiensis, Anopheles stephensi*, is a unique malaria vector because of its ability to thrive in artificial containers in urban environments. This species is found across South and South-East Asia and the Arabian Peninsula, where it is a primary malaria vector and responsible for both urban and rural malaria transmission. Most malaria vector control efforts in Africa are focused on rural habitats, and the ability for malaria vectors to thrive in urban environments may threaten progress made on malaria control and elimination.

In 2012, *An. stephensi* was first detected on the African continent in a livestock quarantine station in a seaport in Djibouti (Faulde et al., 2014). By 2016, it was then detected in neighboring Ethiopia (Carter et al., 2018). By 2018 (Ahmed et al., 2021) or 2019 (World Health Organization, 2019), *An. stephensi* was detected near seaports in Sudan, as well as Somalia in 2019 (World Health Organization, 2019). With *An. stephensi* having unique ecological characteristics and the first detection of the species in seaports, it has been hypothesized that *An. stephensi* introduction was likely facilitated through maritime trade. Further supporting the similarities between *An. stephensi* and *Ae. aegypti* is the fact that in Ethiopia, a large percentage (40%-Balkew et al. or greater -PMI VL 2021) of the habitats where *An. stephensi* larvae were detected, *Ae. aegypti* were also detected. With invasive *An. stephensi* populations now established in these countries, there is a new threat to malaria control on the African continent. Population genetic analyses suggest the potential source of introduction is South Asia (pre-print by Carter et al. 2021).

The invasion of this malaria vector has the potential to significantly impact global malaria control and elimination efforts (Hamlet et al., 2021). For example, in Djibouti, *An. stephensi* was linked to malaria outbreaks in 2013 (Faulde et al., 2014) and since its initial detection in Djibouti, malaria cases have increased 30-fold (World Health Organization, 2020). Additionally, although it shows a seasonal variability in abundance in Asia, it has been detected year-round through the hot, dry season in Africa (Seyfarth et al., 2019). Recent laboratory studies on invasive Djiboutian and Ethiopian *An. stephensi* specimens reveal that, as in Asia, these populations are competent vectors for both *Plasmodium vivax* and *Plasmodium falciparum* (Seyfarth, 2019). Thus, countries may need to expand their malaria testing protocol and use of Rapid Diagnostic Tests (RDTs) to detect *P. vivax* in countries where it is less common. Further, field data have shown confirmation of *P. vivax* sporozoites in *An. stephensi* in Ethiopia (Tadesse et al., 2021), and high levels of resistance to nearly all insecticides used in malaria vector control (Yared et al., 2020).

Primary malaria vector species are found at lower densities in the urban centers of sub-Saharan Africa, and consequently, these cities tend to have lower malaria transmission rates than surrounding rural areas (Robert et al., 2003). However, a recent habitat suitability modeling study predicted that the further invasion of *An. stephensi* into urban locations on the African continent could put an additional 126 million people at risk of malaria (Sinka et al., 2020).

To address this global challenge and proactively mitigate the threat of *An. stephensi*, prioritization activities are necessary to identify where this invasive mosquito is likely to be introduced, particularly if this is facilitated by the movement of cargo through marine shipping. To better understand the potential invasion dynamics of *An. stephensi*, we use United Nations Commerce and Trade (UNCTAD) global data on maritime cargo shipment, including number of shipments, transshipments, cargo volume, and bilateral connectivity, between countries in South Asia and the Arabian Peninsula where *An. stephensi* is endemic and locations on the African continent where *An. stephensi* is invasive, as well as all coastal African nations where *An. stephensi* has not yet been detected. To account for time periods before and after invasion in Africa, data from 2011, 2016, and 2020 were used to identify coastal nations with the highest risk of *An. stephensi* introduction. These data were then combined with an *An. stephensi* habitat suitability ranking developed by Sinka et al. (2020).

In this manuscript we describe: 1) data from 2011, prior to the detection of *An. stephensi* in Djibouti, to determine whether historical maritime connectivity identify Djibouti and Sudan as high risk countries for *An. stephensi* introduction; 2) a prioritized list of coastal African countries for immediate surveillance based on 2020 data to allow for early detection, rapid response, and limit further introduction of the vector in Africa; and 3) an interactive network model of intracontinental transport routes in Africa allowing for future prioritization hierarchies for surveillance if/when *An. stephensi* is detected in new locations.

## Materials and methods

### Days at sea, habitat suitability index, trade index

Due to the initial detection of *An. stephensi* in the port city of Djibouti City, maritime trade data were examined. We ranked the maritime trade connection between countries with known *An. stephensi* populations (India, Pakistan, Saudi Arabia, and United Arab Emirates) and coastal African countries. Other countries with *An. stephensi* populations such as Iraq, Iran, and Thailand, exhibited lower trade levels and were not included. Additionally, Ethiopia and Somalia were not included despite having confirmed *An. stephensi* populations due to the absence of UNCTAD maritime traffic data.

We used UNCTAD’s Liner Shipping Bilateral Connectivity Index (LSBCI), an index created from trade data from MDS Transmodal (https://www.mdst.co.uk), to measure the amount of connectivity between each pair of countries. The LSBCI factors in five maritime trade indicators. The first is the number of transshipments, when goods are unloaded and moved to another vessel, to get from country *j* to country *k*. Secondly, LSBCI factors in the number of countries which have direct routes to both countries in the pair (e.g. four countries have direct connections to both country *j* and country *k*). The third indicator is the number of common connections with one transshipment shared between the countries. The level of competition on services that connect the countries, measured by the number of carriers operating on this route, serves as another indicator. Finally, the size of the largest ship on the route with the fewest carriers is considered in calculating LSBCI for a country pair, which can serve as a metric of capacity on sea routes. Each indicator is normalized by subtracting the minimum value from the raw value and dividing by the range. LSBCI is the simple average of the normalized value of these five indicators (Fugazza & Hoffmann, 2017). This data does not factor in type of vessel or goods, which could influence *An. stephensi* survivability on board.

We took the LSBCI value and divided it by the number of days required to travel by shipping vessel between the closest and largest ports of the countries. This was calculated via *Searoutes* which uses the automatic identification system (AIS) of vessels to track them and calculate average time between ports (Searoutes – Making Supply Chains Greener., n.d.). The same vessel speed was used in this calculation to maintain uniformity in measuring distance. This compiled index includes 1) maritime trade degree of connectivity and 2) time between ports (in days) is referred to as the likelihood of *An. stephensi* introduction through maritime trade index (LASIMTI).

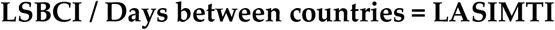

Additionally, we incorporated Sinka et al.’s Habitat Suitability Index (HSI), which uses, in order of importance, annual mean temperature, population density, seasonal precipitation, surface wetness, vegetation, and other environmental factors to evaluate locations with suitable environments for *An. stephensi* establishment. Using R (https://www.r-project.org/), a data set of countries was ranked by LASIMTI as well as both LASIMTI and the HSI.

The UNCTAD trade data from three years - 2011, 2016, and 2020 - were chosen. The year 2011 was selected because it was one year prior to the first detection of *An. stephensi* in the Horn of Africa in Djibouti City. In 2016, *An. stephensi* was confirmed in Ethiopia, potentially indicating further intracontinental spread or separate introductions. However, Ethiopia is landlocked and therefore was not included in this study. Finally, maritime trade data from 2020 was evaluated to assess further spread along this pathway. Potentially important to note, the UNCTAD estimated that maritime trade fell by 4.1% in 2020 due to the COVID-19 pandemic. However, they also predicted a rebound of 4.8% in 2021 (United Nations Conference on Trade and Development, 2021). Additionally, the various types of vessels and goods could influence *An. stephensi* survivability on board.

Maritime trade data from 2020 was used to create a network model of intracontinental African trade between coastal African countries (Figure 3). The connectivity of coastal African nations was examined based on country pairs’ LSBCI. The top three countries, as ranked by LSBCI for each country, were highlighted as links between the nodes. In cases of ties, both countries were included (e.g. Sudan has four country pairs because Egypt, Kenya, and Morocco had the same LSBCI). Another network model was created with a cutoff of 14 days of travel between each node to factor in survivability of *An. stephensi* during transit (Supplemental Figure 1), assuming *An. stephensi* are traveling as eggs which hatch upon arrival into a port based on previous literature describing *An. stephensi* egg resistance to desiccation for nearly two weeks (Chalam 1927). Edges are weighted by the LSBCI value and nodes are weighted by the number of connected countries. Djibouti and Sudan are differentiated due to their established *An. stephensi* populations. This network model was created with R in RStudio utilizing the igraph and visNetwork packages.

**Figure 1.**
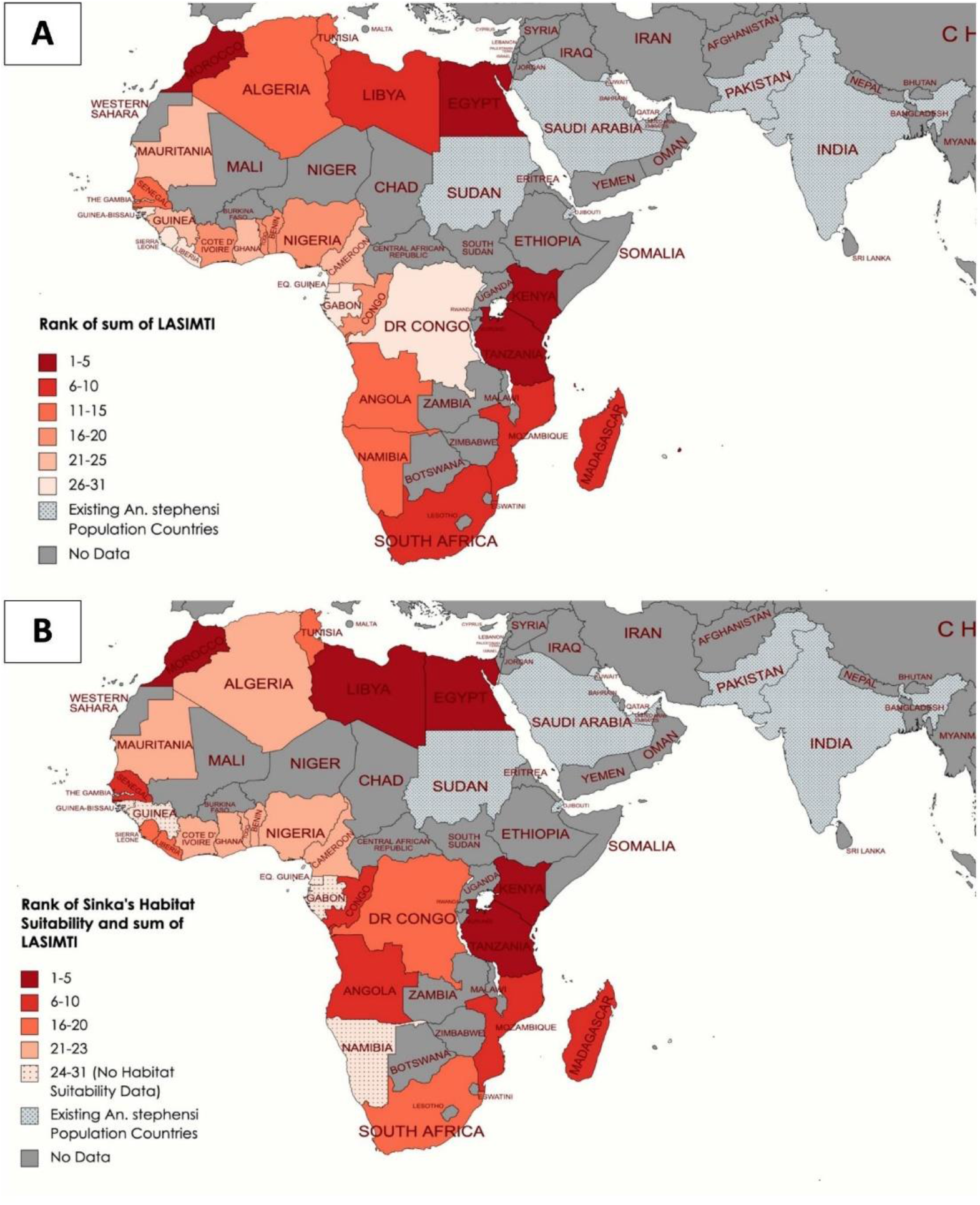
These 2020 heat maps rank coastal African countries using (A) LASIMTI data alone and (B) LASIMTI and HSI combined, based on maritime connectivity to countries where *An. stephensi* is endemic. Higher ranking countries which are at greater risk of *An. stephensi* introduction are darker in red color than those that are lower ranking (lighter red). Countries which are shaded grey are inland countries that do not have a coast or there is no data on maritime movement available. Of those countries without data, Ethiopia and Somalia have confirmed *An. stephensi* populations. Countries which are grey and patterned have established or endemic *An. stephensi* populations and are considered to be source locations for potential *An. stephensi* introduction in this analysis.

**Figure 2.**
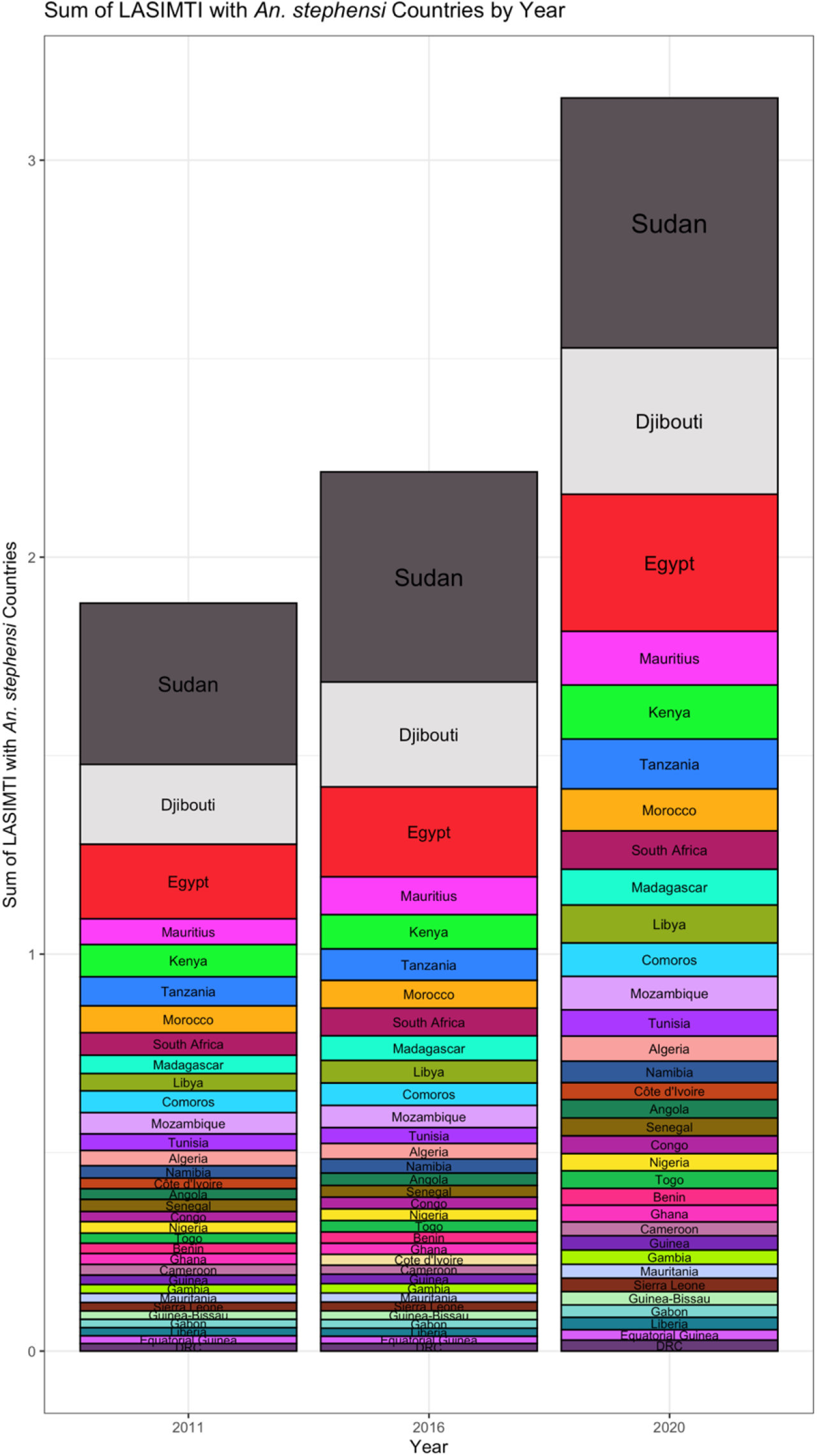
The sum of each LASIMTI for coastal African countries with inputs from endemic *An. stephensi* countries sorted in descending order and arranged by year to highlight highly connected countries and overall maritime traffic growth. This graph breaks down the LASIMTI ranking by country by year. Each column is sorted by count LASIMTI sum. This shows that overall maritime trade between endemic *An. stephensi* countries and coastal Africa has increased over time. This also highlights Sudan, Djibouti, Egypt, Mauritius, Kenya, and Tanzania as highly connected countries.

**Figure 3.**
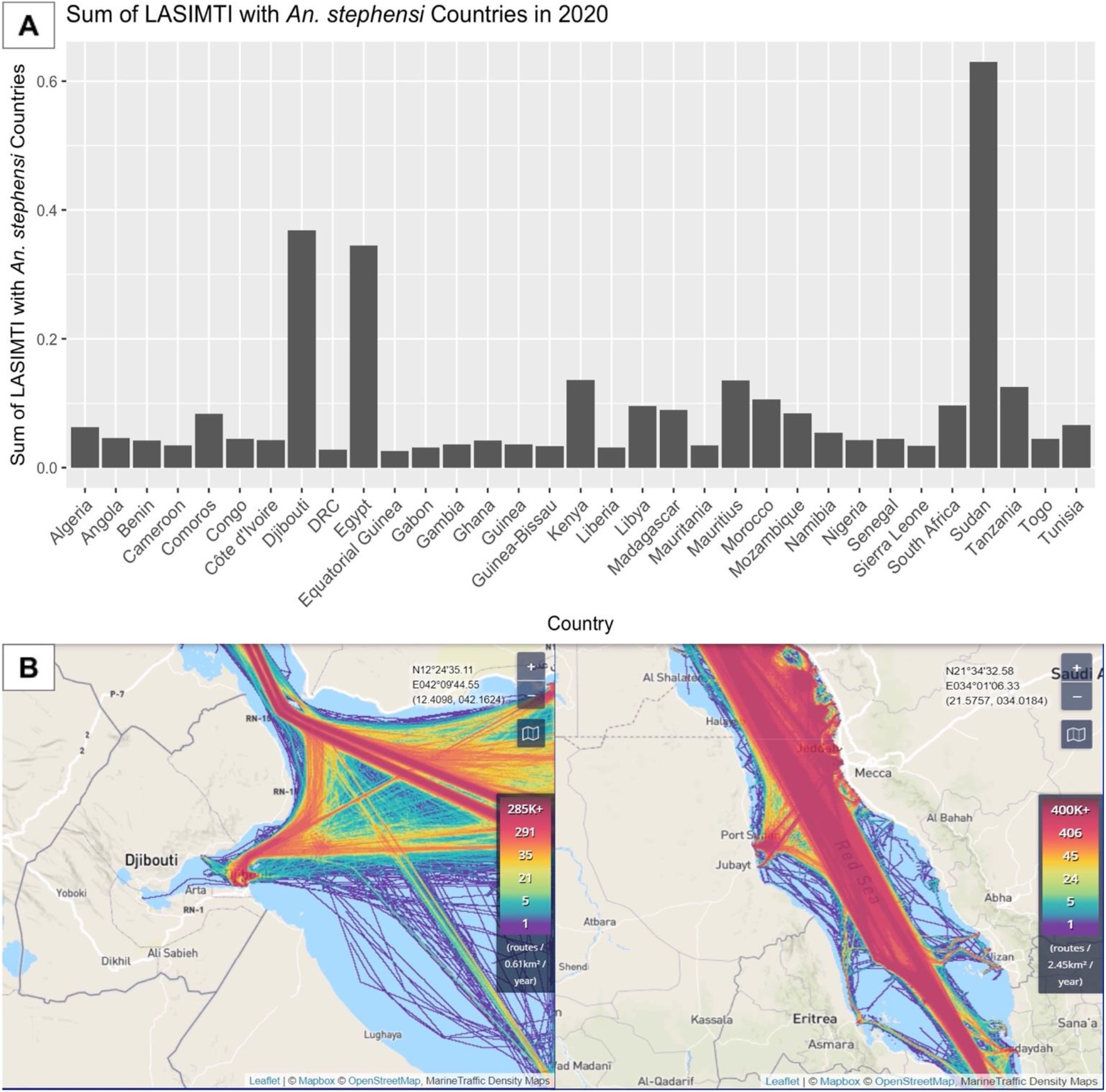
LASIMTI of coastal African countries in 2020 shows heterogeneity across the continent in maritime movement into ports. A) Relatively high traffic from countries where *An. stephensi* is endemic to Egypt, Djibouti, and Sudan. B) Visualization of the volume of traffic into Djibouti and Sudan in 2019 (modified from marinetraffic.com) shows that a few ports in these two countries accommodate hundreds of thousands of transport routes each year.

**Figure 4.**
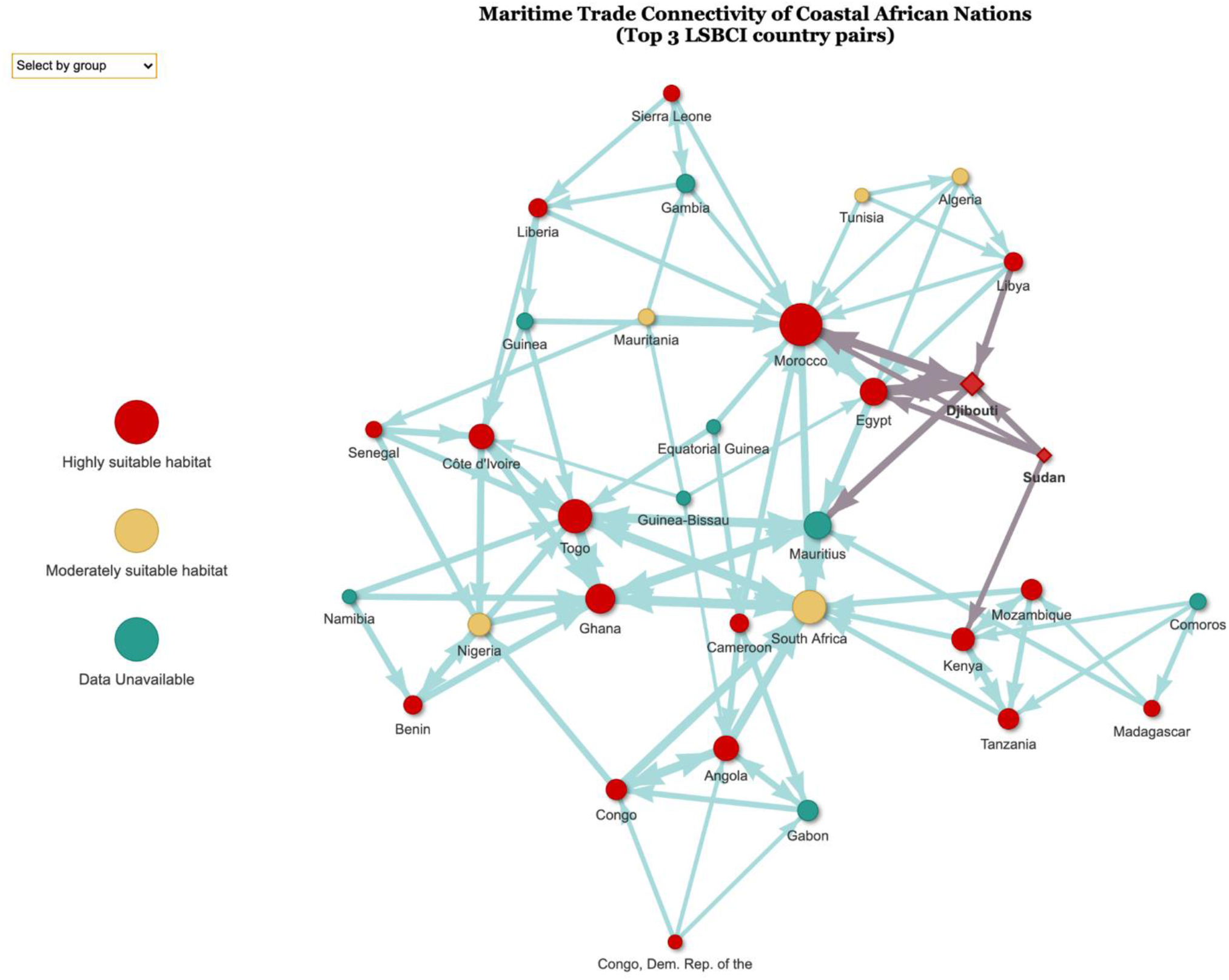
Directed network model of coastal African nations connected through ranking LSBCI data with Sudan and Djibouti highlighted as having known *An. stephensi* populations. This network model was produced using the 2020 UNCTAD trade index, LSBCI. Each node represents a coastal African country with directed edges pointing towards another node. A connection indicates an LSBCI ranked as one of the origin node’s highest three LSBCI. The nodes are also weighted by the number of connections directed towards it as shown by the size. The red diamond nodes (Djibouti and Sudan) are countries with known *An. stephensi* populations. (Interactive HTML link found in supplement)

Network centrality is often calculated with eigenvector centrality, which measures the influence of nodes by factoring in the number of connections and the number of connections of its neighbors. PageRank is a variant of eigenvector centrality but considers the direction of edges. PageRank was used for this network model because of the directed, weighted edges. This rank value determines the centrality of a single node in a network based upon how many connections point towards and away from the node as well as each of its neighbors’ total number of connections. Edge weights and values of other nodes are factored in as well. The PageRank value is ultimately a probability distribution of the nodes in the network. In this network, this would be if a single vessel was selected, the probability that it would be found at a given node. PageRank was calculated in RStudio with the igraph package.

## Results

### *Maritime index in 2011 prior to detection of* Anopheles stephensi *in Africa identified Sudan and Djibouti as highest for risk of introduction*

The 2011 Maritime trade data from UNCTAD pointed to Sudan and Djibouti as the top two connected countries to *An. stephensi* source populations (India, Pakistan, Saudi Arabia, and UAE) when the LASIMTI was summed. The next three countries were Egypt, Kenya, and Tanzania (Table 1, full table: Supplemental Table 1). When HSI was included, the top five countries remained the same (Supplemental Table 2).

**Table 1.** Top 10 coastal countries ranked by *LASIMTI* maritime traffic alone from 2011 UNCTAD maritime trade data.

| Rank | African Country | Sum of LASIMTI |
| --- | --- | --- |
| 1 | Sudan | 0.406 |
| 2 | Djibouti | 0.201 |
| 3 | Egypt | 0.188 |
| 4 | Kenya | 0.081 |
| 5 | Tanzania | 0.073 |
| 6 | Morocco | 0.068 |
| 7 | Mauritius | 0.065 |
| 8 | South Africa | 0.057 |
| 9 | Comoros | 0.055 |
| 10 | Mozambique | 0.054 |

### *Maritime index in 2016 following detection of* Anopheles stephensi *in Djibouti and Ethiopia highlighted Sudan at highest for risk of introduction*

The 2016 UNCTAD maritime trade data shown in Supplemental Table 3 highlighted, in order, Sudan, Djibouti, Egypt, Mauritius, and Kenya when ranked by the sum of LASIMTI to the source *An. stephensi* populations. When this data was ranked first by HSI, the top 5 countries were Sudan, Djibouti, Egypt, Kenya, and Tanzania (Supplemental Table 4)

*Anopheles stephensi* was established in Djibouti in 2012, so after this date, Djibouti was included as a source population in the calculation, which gave the top five countries as Sudan, Egypt, Mauritius, Kenya, and Tanzania when ranked by their sum of LASIMTI to each source population (Supplemental Table 5). The top five countries when ranked by HSI and then LASIMTI sum were Sudan, Egypt, Kenya, Tanzania, and Morocco when Djibouti was included as a source population (Supplemental Table 6).

### *Maritime index in 2020 following detection of* Anopheles stephensi *in Djibouti, Ethiopia, Somalia and Sudan highlighted Kenya, Tanzania, and Mauritius at highest risk of introduction*

The 2020 version of these data indicated Sudan, Djibouti, Egypt, Kenya, and Mauritius as the top five connected countries when ranked by the sum of LASIMTI (Supplemental Table 7). Sudan and Djibouti remained the top two connected countries for all the three years examined. The data utilizing both the HSI and LASIMTI placed Sudan, Djibouti, Egypt, Kenya, and Tanzania as the top five countries (Supplemental Table 8).

Since *An. stephensi* populations have been confirmed in Sudan in 2018 or 2019, these data were further examined with Djibouti and Sudan included as potential source populations for *An. stephensi* (WHO, 2019). With Djibouti and Sudan included as source populations while calculating LASIMTI, the top five countries at risk of *An. stephensi* introduction were Egypt, Kenya, Mauritius, Tanzania, and Morocco (Table 2A). When the HSI was also included in the ordering, the top five countries were Egypt, Kenya, Tanzania, Morocco, and Libya (Table 2B). Full tables can be found in the supplemental material (Supplemental Tables 9 and 10, respectively)

**Table 2.**
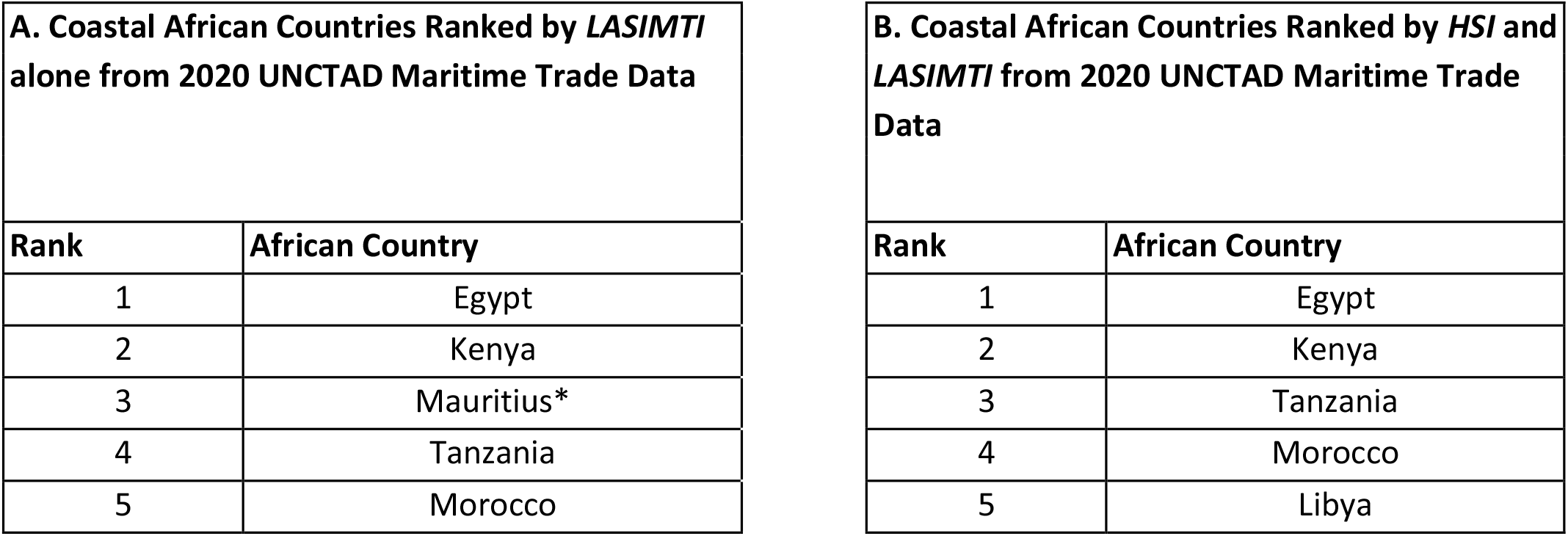
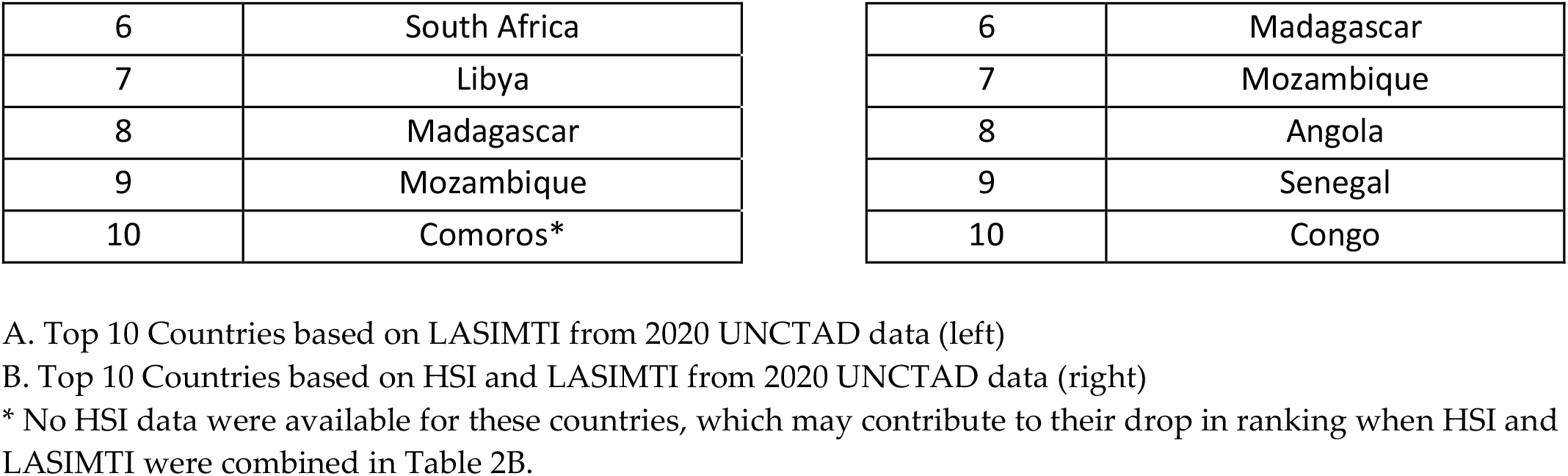
Top 10 coastal countries ranked by *LASIMTI* maritime traffic alone and *LASIMTI* and *HSI* combined using 2020 UNCTAD maritime trade data.

### Intracontinental connectivity network model

The interactive network model reveals degrees of connectivity within coastal nations on the African continent. Specifically, it highlights highly connected coastal African countries such as South Africa as well as the Western African nations. Utilizing the PageRank centrality score, South Africa (0.175), Mauritius (0.159), Ghana (0.159), Togo (0.157), and Morocco (0.044) were more highly connected to coastal countries in Africa than others via maritime trade in this network (Supplemental Table 11). Djibouti and Sudan were ranked 7th (0.030) and 32nd (0.0045) respectively. Egypt was highlighted often as being at risk of introduction by the LASIMTI ranking. In the PageRank centrality analysis, Egypt was ranked 6th with a rank value of 0.0353. Other countries that were highlighted are Kenya (11th, 0.0164) and Tanzania (12th, 0.0156).

## Discussion

With human movement and globalization, invasive container breeding mosquitoes capable of transmitting dengue, Zika, chikungunya and now malaria, with *An. stephensi*, are being introduced and establishing populations in new locations. They are bringing with them the threat of increased risk of vector-borne diseases to new locations where health systems may not be prepared.

*Anopheles stephensi* was first detected on the African continent in Djibouti in 2012 and has since been confirmed in Ethiopia, Somalia, and Sudan. Unlike most malaria vectors, *An. stephensi* is often found in artificial containers and in urban settings. Additionally, historical reports show that *An. stephensi* eggs are able to resist desiccation in soil for up to 14 days (Chalam, 1927). This unique ecology combined with its initial detection in seaports in Djibouti, Somalia, and Sudan has led scientists to believe that the movement of this vector is likely facilitated through maritime trade.

By modeling inter- and intra-continental maritime connectivity in Africa, we identified a ranking of likelihood of *An. stephensi* introduction if facilitated through maritime movement. *Anopheles stephensi* was not detected in Africa (Djibouti) until 2012. To determine whether historical maritime data would have identified the first sites of introduction, 2011 maritime data were analyzed to determine whether the sites with confirmed *An. stephensi* would rank highly in connectivity to *An. stephensi* endemic countries. Using 2011 data on maritime connectivity alone, Djibouti and Sudan were identified as the top two countries at risk of *An. stephensi* introduction facilitated by marine cargo shipments. In 2021, these are two of the three African coastal nations where *An. stephensi* is confirmed to be established.

When 2011 maritime data were combined with the HSI for *An. stephensi* establishment, the top five countries remained the same as with maritime data alone: Sudan, Djibouti, Egypt, Kenya and Tanzania. The maritime data showed the likelihood of introduction, and HSI showed the likelihood of establishment. When combined, the analyses showed a likelihood of being able to establish and survive once introduced. Interestingly, the results of the combined analyses aligned with the detection data being reported in the Horn of Africa. The 2011 maritime data reinforced the validity of the model as it pointed to Sudan and Djibouti as highest risk, and those were the two countries where *An. stephensi* initially established in the following years. Similarly, the HSI data for Ethiopia aligned closely with detections of the species to date (Balkew et al. 2021). Interestingly, around this time of initial detection in Djibouti, Djibouti City port underwent development and organizational change. The government of Djibouti took back administrative control of the port as early as 2012 due disputes with the previous port controller (*Port History – PORT DE DJIBOUTI*, n.d.). The development and construction could have been created environments for the establishment of *An. stephensi*. Additionally, the change of hand might have made it easier for this to go unnoticed.

Following this method, the maritime trade data from 2020 highlighted countries at risk of *An. stephensi* introduction from endemic countries as well as from the coastal African countries with newly introduced populations. We provided a prioritized list of countries for the early detection, rapid response, and targeted surveillance of *An. stephensi* in Africa based on these data and the HSI. Further invasion of *An. stephensi* on the African continent has the potential to reverse progress made on malaria control in the last century and divert resources from rural to urban settings which could worsen the situation. *Anopheles stephensi* thrives in urban settings and in containers, in contrast to the rural settings and natural habitats where most *Anopheles* spp. are found (Seyfarth, 2019). The situation in Djibouti may be a harbinger for what is to come if immediate surveillance and control strategies are not initiated in the countries identified at highest risk (Hamlet et al., 2021).

Maritime data from 2020, with Djibouti and Sudan considered as potential source populations for intracontinental introduction of *An. stephensi*, indicated the top five vulnerable countries were Egypt, Kenya, Mauritius, Tanzania, and Morocco. Targeted larval surveillance in these countries near seaports may provide a better understanding of whether there are maritime introductions. When the 2020 maritime data was combined with HSI for *An. stephensi*, the top five countries were instead Egypt, Kenya, Tanzania, Morocco, and Libya. Interestingly, historical reports of *An. stephensi* in Egypt exist; however, following further identification, these specimens were determined to be *An. ainshamsi* (Gad et al. 2006). With several suitable habitats both along the Egyptian coast and inland, revisiting surveillance efforts there would provide insight into how countries that are highly connected to *An. stephensi*-endemic locations through maritime traffic may experience introductions.

Further field validation of this prioritized list is necessary, because it is possible that *An. stephensi* is being introduced through other transportation routes, such as land transport hubs or airports (Tatem et al. 2006), or may even be dispersed through wind facilitation (Huestis et al., 2019). However, countries highlighted here with high levels of connectivity to known *An. stephensi* locations should be considered at high risk of introduction. Vector and malaria case surveillance should be urgently established to determine whether *An. stephensi* introduction has already occurred. Primary surveillance for both *Aedes aegypti* and *An. stephensi* are through larval surveys, and the two mosquitoes are commonly detected in the same larval habitats. It could therefore be beneficial to coordinate with existing *Aedes* surveillance efforts to be able to simultaneously gather data on medically relevant *Aedes* vectors while seeking to determine whether *An. stephensi* is present. Similarly, in locations with known *An. stephensi* without well-established *Aedes* programs, coordinating larval surveillance efforts would provide an opportunity to conduct malaria and arboviral vector surveillance simultaneously.

Efforts to map key points of introduction based on the movement of goods and people could provide high specificity for targeted surveillance and control efforts. For example, participatory mapping or population mobility data collection methods, such as those used to determine routes of human movement for malaria elimination, may simultaneously provide information on where targeted *An. stephensi* surveillance efforts should focus. Several methods have been proposed in the literature for modeling human movement and one in particular, PopCAB, which is often used for communicable diseases, combined quantitative and qualitative data with geospatial information to identify points of control (Merrill et al., 2021).

Data on invasive mosquito species have shown that introduction events are rarely a one-time occurrence. Population genetics data on *Aedes* species indicate that reintroductions are very common and can facilitate the movement of genes between geographically distinct populations, raising the potential for introduction of insecticide resistance, thermotolerance, and other phenotypic and even behavioral traits which may be facilitated by gene flow and introgression (Yared et al. 2020). Djibouti, Sudan, Somalia, and Ethiopia, countries with established invasive populations of *An. stephensi*, should continue to monitor invasive populations and points of introduction to control and limit further expansion and adaptation of *An. stephensi*. Work by Carter et al. 2021 (pre-print) has shown that *An. stephensi* populations in Ethiopia in the north and central regions can be differentiated genetically, potentially indicating that these populations are a result of more than one introduction into Ethiopia from South Asia, further emphasizing the potential role of anthropogenic movement on the introduction of the species (Carter et al. 2021).

One major limitation of this work is that Somalia is the third coastal nation where *An. stephensi* has been confirmed; however, marine traffic data were not available for Somalia so it could not be included in this analysis. The potential impact of Somalia on maritime trade is unknown and it should not be excluded as a potential source population. Other countries with *An. stephensi* populations, such as Iran, Myanmar, and Iraq, constitute lower relative percentages of trade with these countries so were not included in the analysis. However, genetic similarities were noted between *An. stephensi* in Ethiopia and Pakistan, so this nation was included (Carter et al., 2018).

Due to the nature of maritime traffic, inland countries, such as Ethiopia, were also not included in this prioritized ranking. Countries which are inland but share borders with high-risk countries according to the LASTIMI index should also be considered with high priority. For example, the ranking from 2011 highlights Sudan and Djibouti, both which border Ethiopia, and efforts to examine key international land transportation routes may provide additional insight into the expansion routes of this invasive species.

In Ethiopia, *An. stephensi* was detected in 2016. It has largely been detected along major transportation routes, although further data are needed to understand this association since most sampling sites have also been located along major transport routes. Importantly, Ethiopia relies heavily on the ports of Djibouti and Somalia for maritime imports and exports. Surveillance efforts have revealed that *An. stephensi* is also frequently associated with livestock shelters and are frequently found with livestock bloodmeals (Balkew et al., 2021). The original detection of *An. stephensi* was found in a livestock quarantine station in the port of Djibouti. Additionally, livestock constitutes one of the largest exports of maritime trade from this region. For countries with high maritime connectivity to *An. stephensi* locations but where *An. stephensi* has not been detected, surveillance efforts near seaports, in particular those with livestock trade, is encouraged for early detection of this invasive species.

While not explicitly analyzed in this study, future examination of the movement of specific goods would be beneficial in interpreting potential *An. stephensi* invasion pathways. As *Ae. aegypti* and *Culex coronator* were detected in tires or *Ae. albopictus* through the tire and bamboo (*Dracaena sanderiana)* trade, *An. stephensi* could be carried through maritime trade of specific goods (Higa et al., 2010, Scholte et al., 2008, Yee et al., 2012). Petroleum and oil constitute the largest portion of imports to African nations with established *An. stephensi* populations from the Arabian Peninsula. The second largest group of imports are cars and car parts which could include tires. These African countries primarily exported livestock, coffee, and other agricultural goods (Simoes & Hidalgo, 2011). From the source populations in South Asia, imports to the four African countries with confirmed *Anopheles stephensi* were mainly sugar, textiles, and medicines. Car-related goods including tires were also fairly common imports to these African countries. Exports to South Asian countries consisted of seeds and other agricultural goods as well as crude petroleum. Among the top countries highlighted by the LASIMTI ranking, Djibouti and Sudan imported relatively more sugar and sugar confectionery than Kenya and Tanzania from source population countries in 2019. In 2011, prior to detection of *An. stephensi*, sugar was the highest reported import to Djibouti from these countries.

Egypt, one of the countries that was highlighted in the LASIMTI ranking, imports almost entirely livestock and seeds from the four African countries with known invasive *An. stephensi* populations and exports petroleum, chemical products, and vegetable products. From these same countries, Tanzania and Kenya import large amounts of vehicles while vegetable products also make up a large proportion Kenya’s imports. Mauritius imports mainly precious stones and coffee from the African countries with *An. stephensi*.

Various types of vessels are used to transport certain cargo such as container, bulk, and livestock ships. These vessels could affect *An. stephensi* survivability during transit. Sugar and grain are often shipped in bulk or break bulk vessels which store cargo in large unpackaged containers. Container ships transport products stored in containers sized for land transportation via trucks and carry goods such as tires (UN Conference on Trade and Development, 2018). Livestock vessels are often multilevel, open-air ships which require more hands working on deck and water management (*How is Livestock Transportation Done Using Livestock Carriers?*, 2019).

Using LSBCI index data from 2020, we developed a network to highlight how coastal African nations are connected to through maritime trade. The role of this network analysis was two-fold, 1) it demonstrated intracontinental maritime connectivity; and 2) it highlighted the top three countries connected via maritime trade through an interactive HTML model (Supplemental File). For example, if *An. stephensi* is detected and established in a specific coastal African nation such as Djibouti, selecting the Djibouti node reveals the top three locations at risk of introduction from that source country (Djibouti links to Sudan, Egypt and Kenya). This can be used as an actionable prioritized list for surveillance if *An. stephensi* is detected in any given country and highlights major maritime hubs in Africa which could be targeted for surveillance and control.

The network analysis revealed the significance of South African trade to the rest of the continent. Due to the distance, South Africa did not appear to be at high risk of *An. stephensi* introduction. However, this analysis revealed that if *An. stephensi* were to enter nearby countries, it could very easily be introduced because of its high centrality. Western African countries such as Ghana, Togo, and Morocco are also heavily connected to other parts of Africa. Interestingly, Mauritius appears to be highly significant to this network of African maritime trade. Based on 2020 maritime data, Mauritius was ranked as the country with the third greatest likelihood of introduction of *An. stephensi* and also had the second highest centrality rank value of 0.159. Considering these factors, Mauritius could serve as an important port of call connecting larger ports throughout Africa or other continents. If *An. stephensi* were to become established in countries with high centrality ranks, further expansion on the continent could be accelerated drastically. These ports could serve as important watchpoints and indicators of *An. stephensi*’s incursion into Africa.

*Anopheles stephensi* is often found in shared habitats with *Aedes* spp. and an important opportunity exists to leverage *Aedes* surveillance efforts to detect invasive *An. stephensi*, especially in countries that have high potential of introduction through maritime trade. For example, the island of Mauritius ranks third most connected to *An. stephensi* locations based on 2020 maritime data. With long standing regular larval surveillance efforts across the island for *Aedes* spp., this island nation is well suited to look for *Anopheles* larvae as part of *Aedes* surveillance efforts for early detection and rapid response to prevent the establishment of *An. stephensi*.

## Conclusions

With increases in globalization and the volume and frequency of marine cargo traffic connecting countries and continents, information on maritime connectivity can serve as an early warning system for invasive species in general, including those relevant to public health. We show that maritime data prior to the detection of *An. stephensi* in Africa identified Djibouti and Sudan as countries at greatest risk of introduction, and these are locations where invasive *An. stephensi* populations are now established. Using more recent data we present a prioritized list of countries at risk of *An. stephensi* introduction through maritime traffic and describe intracontinental maritime connectivity. These data highlight the potential use of maritime trade data to guide intensified surveillance efforts for the early detection of invasive mosquito vectors, such as *An. stephensi* in Africa, to limit establishment and impact on public health.

Through integrated vector management, existing *Aedes* programs could be leveraged by providing training for *An. stephensi* identification. The distinct larval characteristics of *Ae. aegypti* and *An. stephensi* suggest that as part of *Aedes* surveillance, the simple addition of also looking for *An. stephensi* larvae can provide opportunities to search for the invasive malaria vector without needing to establish new programs. Similarly, in locations where *An. stephensi* surveillance is ongoing, the addition of data collection on *Aedes* spp. could also be included. These integrated efforts will strengthen local, regional, and national entomological surveillance systems for vector borne diseases.

## Supporting information

Supplementary interactive network model

Supplementary interactive network model 14 day cut off

## Abbreviations

LSBCI: Liner Shipping Bilateral Connectivity Index
LASIMTI: Likelihood of *Anopheles stephensi* Introduction Through Maritime Trade Index
HIS: Habitat suitability index

## Acknowledgements

The authors would like to thank Jen Armistead, Tony Hughes, Audrey Lenhart, John Gimnig, Melissa Yoshimizu, Peter Mumba, and Diana Iyaloo for providing valuable feedback on previous versions of the manuscript and Anne Wilson, Dave Weetman, Ayman Ahmed, and Tamar Carter for fruitful discussions on the approach.

## Consent for publication

Not applicable

## Availability of data and material

All data generated are included in this manuscript and supplementary files.

## Competing interests

The authors declare that they have no competing interests.

## Funding

JA is funded by Emory University and the U.S. Centers for Disease Control and Prevention. SZ and SI are funded by the U.S. President’s Malaria Initiative.

## Disclaimer

The findings and conclusions in this report are those of the author(s) and do not necessarily represent the official position of the U.S. Centers for Disease Control and Prevention or the Department of Health and Human Services.

## Notes

### Competing Interest Statement

The authors have declared no competing interest.

