## Supplementary material for "Using marine cargo traffic to identify countries in Africa with greatest risk of invasion by *Anopheles stephensi*"

### Supplemental Figure 1: Network Model based on LSBCI with ≤ 14 days of travel between

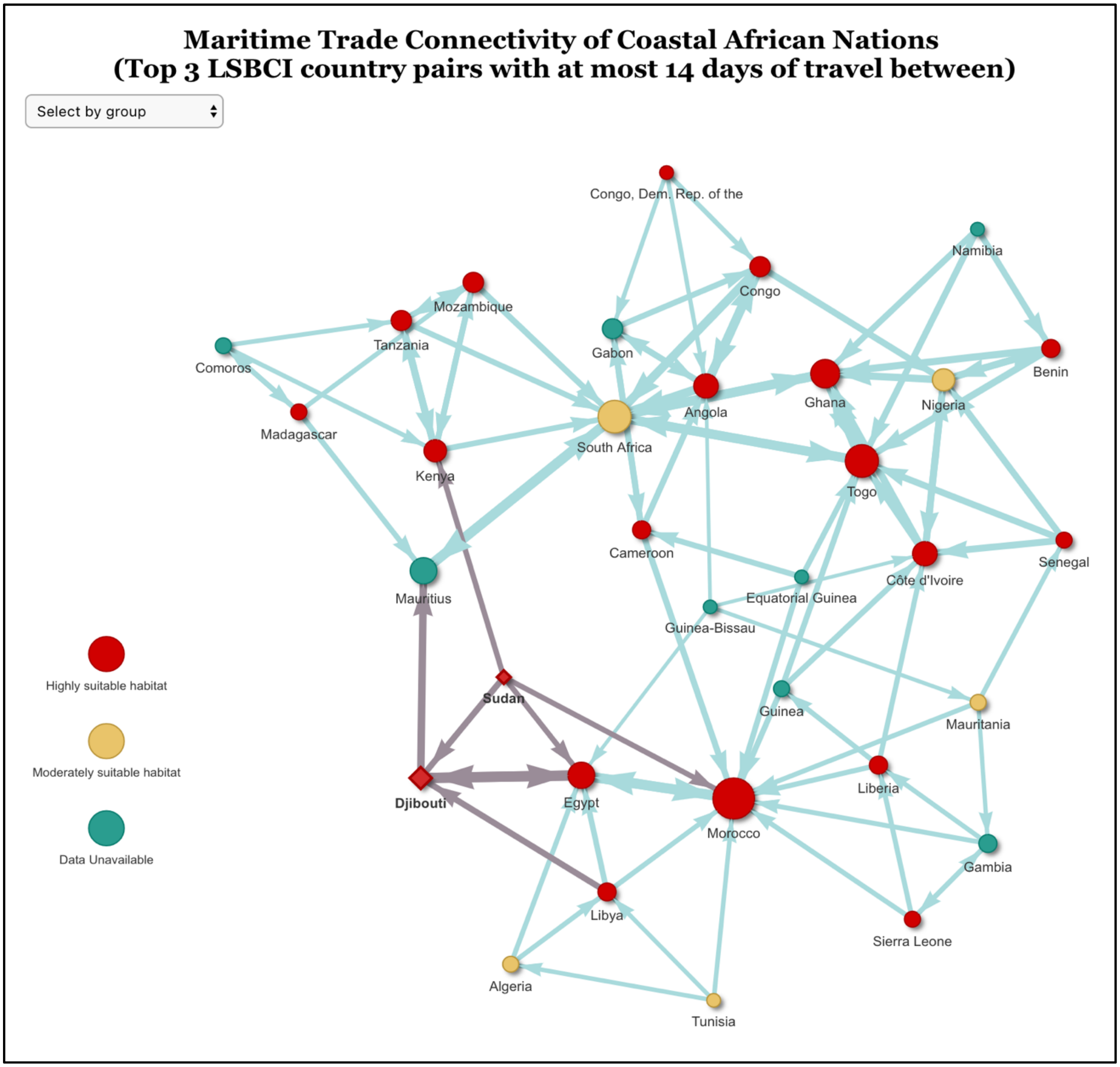


### Supplemental Table 1: 2011 UNCTAD Data- Ranked by Sum of LASIMTI

| Rank | African Country | Sum of LASIMTI | Sinka Habitat Suitability (HSI) |
|---|---|---|---|
| 1 | Sudan | 0.406269375 | 1 |
| 2 | Djibouti | 0.2013548498 | 1 |
| 3 | Egypt | 0.1876630554 | 1 |
| 4 | Kenya | 0.08135093502 | 1 |
| 5 | Tanzania, United Republic of | 0.07330365343 | 1 |
| 6 | Morocco | 0.06757871437 | 1 |
| 7 | Mauritius | 0.064657715 | #N/A |
| 8 | South Africa | 0.05658928249 | 2 |
| 9 | Comoros | 0.05465475623 | #N/A |
| 10 | Mozambique | 0.05389609371 | 1 |
| 11 | Madagascar | 0.04642226481 | 1 |
| 12 | Libya | 0.04345450808 | 1 |
| 13 | Tunisia | 0.04181758212 | 2 |
| 14 | Algeria | 0.03839803059 | 2 |
| 15 | Namibia | 0.03096978557 | #N/A |
| 16 | Senegal | 0.03068857176 | 1 |
| 17 | Nigeria | 0.0286509546 | 2 |
| 18 | Ghana | 0.02753747632 | 1 |
| 19 | Cameroon | 0.02709093522 | 1 |
| 20 | Côte d'Ivoire | 0.02688200356 | 1 |
| 21 | Angola | 0.02667341745 | 1 |
| 22 | Benin | 0.02584286755 | 1 |
| 23 | Togo | 0.02562999872 | 1 |
| 24 | Republic of Congo | 0.02561828986 | 1 |
| 25 | Guinea | 0.02354817407 | #N/A |
| 26 | Mauritania | 0.02318495186 | 2 |
| 27 | Gambia | 0.02204206605 | #N/A |
| 28 | Sierra Leone | 0.02152680573 | 1 |
| 29 | Gabon | 0.02097385197 | #N/A |
| 30 | Liberia | 0.02077916229 | 1 |

| | | | |
|---|---|---|---|
| 31 | Guinea-Bissau | 0.02075651111 | #N/A |
| 32 | Equatorial Guinea | 0.01935484039 | #N/A |
| 33 | Congo, Dem. Rep. of the | 0.01904588403 | 1 |

**Supplemental Table 2: UNCTAD Data – Ranked by HSI and Sum of LASIMTI**

| Rank | African Country | Sum of LASIMTI | HSI |
|---|---|---|---|
| 1 | Sudan | 0.406269375 | 1 |
| 2 | Djibouti | 0.2013548498 | 1 |
| 3 | Egypt | 0.1876630554 | 1 |
| 4 | Kenya | 0.08135093502 | 1 |
| 5 | Tanzania, United Republic of | 0.07330365343 | 1 |
| 6 | Morocco | 0.06757871437 | 1 |
| 7 | Mozambique | 0.05389609371 | 1 |
| 8 | Madagascar | 0.04642226481 | 1 |
| 9 | Libya | 0.04345450808 | 1 |
| 10 | Senegal | 0.03068857176 | 1 |
| 11 | Ghana | 0.02753747632 | 1 |
| 12 | Cameroon | 0.02709093522 | 1 |
| 13 | Côte d'Ivoire | 0.02688200356 | 1 |
| 14 | Angola | 0.02667341745 | 1 |
| 15 | Benin | 0.02584286755 | 1 |
| 16 | Togo | 0.02562999872 | 1 |
| 17 | Republic of Congo | 0.02561828986 | 1 |
| 18 | Sierra Leone | 0.02152680573 | 1 |
| 19 | Liberia | 0.02077916229 | 1 |
| 20 | Congo, Dem. Rep. of the | 0.01904588403 | 1 |
| 21 | South Africa | 0.05658928249 | 2 |
| 22 | Tunisia | 0.04181758212 | 2 |
| 23 | Algeria | 0.03839803059 | 2 |
| 24 | Nigeria | 0.0286509546 | 2 |
| 25 | Mauritania | 0.02318495186 | 2 |
| 26 | Mauritius | 0.064657715 | #N/A |
| 27 | Comoros | 0.05465475623 | #N/A |
| 28 | Namibia | 0.03096978557 | #N/A |

| | | | |
|---|---|---|---|
| 29 | Guinea | 0.02354817407 | #N/A |
| 30 | Gambia | 0.02204206605 | #N/A |
| 31 | Gabon | 0.02097385197 | #N/A |
| 32 | Guinea-Bissau | 0.02075651111 | #N/A |
| 33 | Equatorial Guinea | 0.01935484039 | #N/A |

**Supplemental Table 3: 2016 UNCTAD Data – Ranked by Sum of LASIMTI**

| **Rank** | **African Country** | **Sum of LASIMTI** | **HSI** |
|---|---|---|---|
| 1 | Sudan | 0.5292875582 | 1 |
| 2 | Djibouti | 0.2642345551 | 1 |
| 3 | Egypt | 0.2264942296 | 1 |
| 4 | Mauritius | 0.09515349111 | #N/A |
| 5 | Kenya | 0.086537827 | 1 |
| 6 | Tanzania, United Republic of | 0.07930393022 | 1 |
| 7 | Morocco | 0.06989300335 | 1 |
| 8 | South Africa | 0.069568283 | 2 |
| 9 | Madagascar | 0.06215963634 | 1 |
| 10 | Libya | 0.05686532422 | 1 |
| 11 | Comoros | 0.05628177587 | #N/A |
| 12 | Mozambique | 0.05608540859 | 1 |
| 13 | Tunisia | 0.03995433025 | 2 |
| 14 | Algeria | 0.03901809148 | 2 |
| 15 | Namibia | 0.03544483498 | #N/A |
| 16 | Angola | 0.03122483304 | 1 |
| 17 | Senegal | 0.02959448458 | 1 |
| 18 | Congo | 0.02949198772 | 1 |
| 19 | Nigeria | 0.02914460952 | 2 |
| 20 | Togo | 0.02897381889 | 1 |
| 21 | Benin | 0.02845344977 | 1 |
| 22 | Ghana | 0.0279808366 | 1 |
| 23 | Cote d'Ivoire | 0.0273037052 | 1 |
| 24 | Guinea | 0.0239046287 | #N/A |
| 25 | Gambia | 0.02349013522 | #N/A |
| 26 | Mauritania | 0.02271279802 | 2 |
| 27 | Cameroon | 0.02258063866 | 1 |
| 28 | Sierra Leone | 0.02234961057 | 1 |
| 29 | Guinea-Bissau | 0.02186597058 | #N/A |
| 30 | Gabon | 0.02159819569 | #N/A |

| | | | |
|---|---|---|---|
| 31 | Liberia | 0.02058087751 | 1 |
| 32 | Congo, Dem. Rep. of the | 0.0192339785 | 1 |
| 33 | Equatorial Guinea | 0.01796302182 | #N/A |

**Supplemental Table 4: 2016 UNCTAD Data – Ranked by HSI and Sum of LASIMTI**

| Rank | African Country | Sum of LASIMTI | HSI |
|---|---|---|---|
| 1 | Sudan | 0.5292875582 | 1 |
| 2 | Djibouti | 0.2642345551 | 1 |
| 3 | Egypt | 0.2264942296 | 1 |
| 4 | Kenya | 0.086537827 | 1 |
| 5 | Tanzania, United Republic of | 0.07930393022 | 1 |
| 6 | Morocco | 0.06989300335 | 1 |
| 7 | Madagascar | 0.06215963634 | 1 |
| 8 | Libya | 0.05686532422 | 1 |
| 9 | Mozambique | 0.05608540859 | 1 |
| 10 | Angola | 0.03122483304 | 1 |
| 11 | Senegal | 0.02959448458 | 1 |
| 12 | Congo | 0.02949198772 | 1 |
| 13 | Togo | 0.02897381889 | 1 |
| 14 | Benin | 0.02845344977 | 1 |
| 15 | Ghana | 0.0279808366 | 1 |
| 16 | Cote d'Ivoire | 0.0273037052 | 1 |
| 17 | Cameroon | 0.02258063866 | 1 |
| 18 | Sierra Leone | 0.02234961057 | 1 |
| 19 | Liberia | 0.02058087751 | 1 |
| 20 | Congo, Dem. Rep. of the | 0.0192339785 | 1 |
| 21 | South Africa | 0.069568283 | 2 |
| 22 | Tunisia | 0.03995433025 | 2 |
| 23 | Algeria | 0.03901809148 | 2 |
| 24 | Nigeria | 0.02914460952 | 2 |
| 25 | Mauritania | 0.02271279802 | 2 |
| 26 | Mauritius | 0.09515349111 | #N/A |
| 27 | Comoros | 0.05628177587 | #N/A |
| 28 | Namibia | 0.03544483498 | #N/A |
| 29 | Guinea | 0.0239046287 | #N/A |
| 30 | Gambia | 0.02349013522 | #N/A |
| 31 | Guinea-Bissau | 0.02186597058 | #N/A |
| 32 | Gabon | 0.02159819569 | #N/A |

| | | | |
|---|---|---|---|
| 33 | Equatorial Guinea | 0.01796302182 | #N/A |

**Supplemental Table 5: 2016 UNCTAD Data – Ranked by Sum of LASIMTI including Djibouti**

| Rank | African Country | Sum of LASIMTI | HSI |
|---|---|---|---|
| 1 | Sudan | 0.6297510767 | 1 |
| 2 | Egypt | 0.2834920893 | 1 |
| 3 | Mauritius | 0.1189301382 | #N/A |
| 4 | Kenya | 0.1153847996 | 1 |
| 5 | Tanzania, United Republic of | 0.1061623789 | 1 |
| 6 | Morocco | 0.08913120335 | 1 |
| 7 | South Africa | 0.08436727571 | 2 |
| 8 | Madagascar | 0.07764632706 | 1 |
| 9 | Libya | 0.07601938483 | 1 |
| 10 | Mozambique | 0.07310251478 | 1 |
| 11 | Comoros | 0.07160145822 | #N/A |
| 12 | Tunisia | 0.05223401597 | 2 |
| 13 | Algeria | 0.05050180815 | 2 |
| 14 | Namibia | 0.0451833777 | #N/A |
| 15 | Angola | 0.03928818897 | 1 |
| 16 | Senegal | 0.03786745958 | 1 |
| 17 | Congo | 0.03738704078 | 1 |
| | Togo | 0.0364840035 | 1 |
| 19 | Nigeria | 0.03605223594 | 2 |
| 20 | Benin | 0.03522935472 | 1 |
| 21 | Ghana | 0.0349352077 | 1 |
| 22 | Côte d'Ivoire | 0.03464305886 | 1 |
| 23 | Guinea | 0.0306644075 | #N/A |
| 24 | Gambia | 0.03026011551 | #N/A |
| 25 | Mauritania | 0.02938804933 | 2 |
| 26 | Sierra Leone | 0.02875647358 | 1 |
| 27 | Guinea-Bissau | 0.02822034857 | #N/A |
| 28 | Cameroon | 0.02814167795 | 1 |
| 29 | Gabon | 0.02712714954 | #N/A |
| 30 | Liberia | 0.02661193037 | 1 |
| 31 | Congo, Dem. Rep. of the | 0.02437425605 | 1 |
| 32 | Equatorial Guinea | 0.02259842486 | #N/A |

**Supplemental Table 6: 2016 UNCTAD Data – Ranked by HIS and Sum of LASIMTI including Djibouti**

| Rank | African Country | Sum of LASIMTI | HSI |
|---|---|---|---|
| 1 | Sudan | 0.6297510767 | 1 |
| 2 | Egypt | 0.2834920893 | 1 |
| 3 | Kenya | 0.1153847996 | 1 |
| 4 | Tanzania, United Republic of | 0.1061623789 | 1 |
| 5 | Morocco | 0.08913120335 | 1 |
| 6 | Madagascar | 0.07764632706 | 1 |
| 7 | Libya | 0.07601938483 | 1 |
| 8 | Mozambique | 0.07310251478 | 1 |
| 9 | Angola | 0.03928818897 | 1 |
| 10 | Senegal | 0.03786745958 | 1 |
| 11 | Congo | 0.03738704078 | 1 |
| 12 | Togo | 0.0364840035 | 1 |
| 13 | Benin | 0.03522935472 | 1 |
| 14 | Ghana | 0.0349352077 | 1 |
| 15 | Côte d'Ivoire | 0.03464305886 | 1 |
| 16 | Sierra Leone | 0.02875647358 | 1 |
| 17 | Cameroon | 0.02814167795 | 1 |
| 18 | Liberia | 0.02661193037 | 1 |
| 19 | Congo, Dem. Rep. of the | 0.02437425605 | 1 |
| 20 | South Africa | 0.08436727571 | 2 |
| 21 | Tunisia | 0.05223401597 | 2 |
| 22 | Algeria | 0.05050180815 | 2 |
| 23 | Nigeria | 0.03605223594 | 2 |
| 24 | Mauritania | 0.02938804933 | 2 |
| 25 | Mauritius | 0.1189301382 | #N/A |
| 26 | Comoros | 0.07160145822 | #N/A |
| 27 | Namibia | 0.0451833777 | #N/A |
| 28 | Guinea | 0.0306644075 | #N/A |
| 29 | Gambia | 0.03026011551 | #N/A |
| 30 | Guinea-Bissau | 0.02822034857 | #N/A |
| 31 | Gabon | 0.02712714954 | #N/A |
| 32 | Equatorial Guinea | 0.02259842486 | #N/A |

**Supplemental Table 7: 2020 UNCTAD Data – Ranked by Sum of LASIMTI**

| Rank | African Country | Sum of LASIMTI | HSI |
|---|---|---|---|
| 1 | Sudan | 0.6297510767 | 1 |
| 2 | Djibouti | 0.3685620551 | 1 |
| 3 | Egypt | 0.345237433 | 1 |
| 4 | Kenya | 0.1361832312 | 1 |
| 5 | Mauritius | 0.1352187222 | #N/A |
| 6 | Tanzania, United Republic of | 0.1257252601 | 1 |
| 7 | Morocco | 0.1056325617 | 1 |
| 8 | South Africa | 0.09662923821 | 2 |
| 9 | Libya | 0.09573710105 | 1 |
| 10 | Madagascar | 0.08997633539 | 1 |
| 11 | Mozambique | 0.08463453701 | 1 |
| 12 | Comoros | 0.08393858785 | #N/A |
| 13 | Tunisia | 0.06613352214 | 2 |
| 14 | Algeria | 0.06305019762 | 2 |
| 15 | Namibia | 0.05400225608 | #N/A |
| 16 | Angola | 0.04645958458 | 1 |
| 17 | Senegal | 0.04494589958 | 1 |
| 18 | Congo | 0.04477493134 | 1 |
| 19 | Togo | 0.0445830801 | 1 |
| 20 | Nigeria | 0.04307946927 | 2 |
| 21 | Côte d'Ivoire | 0.04272174664 | 1 |
| 22 | Benin | 0.04227994715 | 1 |
| 23 | Ghana | 0.04214779211 | 1 |
| 24 | Guinea | 0.03622271479 | #N/A |
| 25 | Gambia | 0.03587745259 | #N/A |
| 26 | Mauritania | 0.03475354634 | 2 |
| 27 | Cameroon | 0.03464224265 | 1 |
| 28 | Sierra Leone | 0.03400713338 | 1 |
| 29 | Guinea-Bissau | 0.03360291614 | #N/A |
| 30 | Liberia | 0.03132986141 | 1 |
| 31 | Gabon | 0.03112523189 | #N/A |
| 32 | Congo, Dem. Rep. of the | 0.02775588197 | 1 |
| 33 | Equatorial Guinea | 0.02596818172 | #N/A |

**Supplemental Table 8: 2020 UNCTAD Data – Ranked by HSI and Sum of LASIMTI**

| Rank | African Country | Sum of LASIMTI | HSI |
|---|---|---|---|
| 1 | Sudan | 0.6297510767 | 1 |
| 2 | Djibouti | 0.3685620551 | 1 |
| 3 | Egypt | 0.345237433 | 1 |
| 4 | Kenya | 0.1361832312 | 1 |
| 5 | Tanzania, United Republic of | 0.1257252601 | 1 |
| 6 | Morocco | 0.1056325617 | 1 |
| 7 | Libya | 0.09573710105 | 1 |
| 8 | Madagascar | 0.08997633539 | 1 |
| 9 | Mozambique | 0.08463453701 | 1 |
| 10 | Angola | 0.04645958458 | 1 |
| 11 | Senegal | 0.04494589958 | 1 |
| 12 | Congo | 0.04477493134 | 1 |
| 13 | Togo | 0.0445830801 | 1 |
| 14 | Côte d'Ivoire | 0.04272174664 | 1 |
| 15 | Benin | 0.04227994715 | 1 |
| 16 | Ghana | 0.04214779211 | 1 |
| 17 | Cameroon | 0.03464224265 | 1 |
| 18 | Sierra Leone | 0.03400713338 | 1 |
| 19 | Liberia | 0.03132986141 | 1 |
| 20 | Congo, Dem. Rep. of the | 0.02775588197 | 1 |
| 21 | South Africa | 0.09662923821 | 2 |
| 22 | Tunisia | 0.06613352214 | 2 |
| 23 | Algeria | 0.06305019762 | 2 |
| 24 | Nigeria | 0.04307946927 | 2 |
| 25 | Mauritania | 0.03475354634 | 2 |
| 26 | Mauritius | 0.1352187222 | #N/A |
| 27 | Comoros | 0.08393858785 | #N/A |
| 28 | Namibia | 0.05400225608 | #N/A |
| 29 | Guinea | 0.03622271479 | #N/A |
| 30 | Gambia | 0.03587745259 | #N/A |
| 31 | Guinea-Bissau | 0.03360291614 | #N/A |
| 32 | Gabon | 0.03112523189 | #N/A |
| 33 | Equatorial Guinea | 0.02596818172 | #N/A |

**Supplemental Table 9: 2020 UNCTAD Data – Ranked by Sum of LASIMTI (Djibouti and Sudan as source population)**

| Rank | African Country | Sum of LASIMTI | HSI |
|---|---|---|---|
| 1 | Egypt | 0.345237433 | 1 |
| 2 | Kenya | 0.1361832312 | 1 |
| 3 | Mauritius | 0.1352187222 | #N/A |
| 4 | Tanzania, United Republic of | 0.1257252601 | 1 |
| 5 | Morocco | 0.1056325617 | 1 |
| 6 | South Africa | 0.09662923821 | 2 |
| 7 | Libya | 0.09573710105 | 1 |
| 8 | Madagascar | 0.08997633539 | 1 |
| 9 | Mozambique | 0.08463453701 | 1 |
| 10 | Comoros | 0.08393858785 | #N/A |
| 11 | Tunisia | 0.06613352214 | 2 |
| 12 | Algeria | 0.06305019762 | 2 |
| 13 | Namibia | 0.05400225608 | #N/A |
| 14 | Angola | 0.04645958458 | 1 |
| 15 | Senegal | 0.04494589958 | 1 |
| 16 | Congo | 0.04477493134 | 1 |
|  | Togo | 0.0445830801 | 1 |
| 18 | Nigeria | 0.04307946927 | 2 |
| 19 | Côte d'Ivoire | 0.04272174664 | 1 |
| 20 | Benin | 0.04227994715 | 1 |
| 21 | Ghana | 0.04214779211 | 1 |
| 22 | Guinea | 0.03622271479 | #N/A |
| 23 | Gambia | 0.03587745259 | #N/A |
| 24 | Mauritania | 0.03475354634 | 2 |
| 25 | Cameroon | 0.03464224265 | 1 |
| 26 | Sierra Leone | 0.03400713338 | 1 |
| 27 | Guinea-Bissau | 0.03360291614 | #N/A |
| 28 | Liberia | 0.03132986141 | 1 |
| 29 | Gabon | 0.03112523189 | #N/A |
| 30 | Congo, Dem. Rep. of the | 0.02775588197 | 1 |
| 31 | Equatorial Guinea | 0.02596818172 | #N/A |

**Supplemental Table 10: 2020 UNCTAD Data – ranked by HSI and Sum of LASIMTI (Djibouti and Sudan as source population)**

| Rank | African Country | Sum of LASIMTI | HSI |
|---|---|---|---|
| 1 | Egypt | 0.345237433 | 1 |
| 2 | Kenya | 0.1361832312 | 1 |
| 3 | Tanzania, United Republic of | 0.1257252601 | 1 |
| 4 | Morocco | 0.1056325617 | 1 |
| 5 | Libya | 0.09573710105 | 1 |
| 6 | Madagascar | 0.08997633539 | 1 |
| 7 | Mozambique | 0.08463453701 | 1 |
| 8 | Angola | 0.04645958458 | 1 |
| 9 | Senegal | 0.04494589958 | 1 |
| 10 | Congo | 0.04477493134 | 1 |
| 11 | Togo | 0.0445830801 | 1 |
| 12 | Côte d'Ivoire | 0.04272174664 | 1 |
| 13 | Benin | 0.04227994715 | 1 |
| 14 | Ghana | 0.04214779211 | 1 |
| 15 | Cameroon | 0.03464224265 | 1 |
| 16 | Sierra Leone | 0.03400713338 | 1 |
| 17 | Liberia | 0.03132986141 | 1 |
| 18 | Congo, Dem. Rep. of the | 0.02775588197 | 1 |
| 19 | South Africa | 0.09662923821 | 2 |
| 20 | Tunisia | 0.06613352214 | 2 |
| 21 | Algeria | 0.06305019762 | 2 |
| 22 | Nigeria | 0.04307946927 | 2 |
| 23 | Mauritania | 0.03475354634 | 2 |
| 24 | Mauritius | 0.1352187222 | #N/A |
| 25 | Comoros | 0.08393858785 | #N/A |
| 26 | Namibia | 0.05400225608 | #N/A |
| 27 | Guinea | 0.03622271479 | #N/A |
| 28 | Gambia | 0.03587745259 | #N/A |
| 29 | Guinea-Bissau | 0.03360291614 | #N/A |
| 30 | Gabon | 0.03112523189 | #N/A |
| 31 | Equatorial Guinea | 0.02596818172 | #N/A |

**Supplemental Table 11: PageRank**

| Country | Rank | PageRank value |
|---|---|---|
| South Africa | 1 | 0.174839853 |
| Mauritius | 2 | 0.1591659086 |
| Ghana | 3 | 0.1589446226 |
| Togo | 4 | 0.157321488 |
| Morocco | 5 | 0.04406940752 |
| Egypt | 6 | 0.03533168663 |
| Djibouti | 7 | 0.02959821682 |
| Angola | 8 | 0.01757490797 |
| Nigeria | 9 | 0.0173026753 |
| Côte d'Ivoire | 10 | 0.01649370161 |
| Kenya | 11 | 0.01636531004 |
| Tanzania | 12 | 0.01559391596 |
| Congo | 13 | 0.01490935531 |
| Mozambique | 14 | 0.01472534712 |
| Gabon | 15 | 0.01225321047 |
| Benin | 16 | 0.01067750093 |
| Cameroon | 17 | 0.009371850407 |
| Liberia | 18 | 0.008686871017 |
| Gambia | 19 | 0.007893176808 |
| Libya | 20 | 0.007370625956 |
| Guinea | 21 | 0.006989122645 |
| Sierra Leone | 22 | 0.00679030984 |
| Madagascar | 23 | 0.006474458646 |
| Comoros | 24 | 0.006376567262 |
| Senegal | 25 | 0.006179782093 |
| Algeria | 26 | 0.005835988754 |
| Mauritania | 27 | 0.005591411351 |
| Congo, Dem. Rep. of the | 28 | 0.004545454545 |
| Equatorial Guinea | 29 | 0.004545454545 |
| Guinea-Bissau | 30 | 0.004545454545 |
| Namibia | 31 | 0.004545454545 |
| Sudan | 32 | 0.004545454545 |
| Tunisia | 33 | 0.004545454545 |
